# Behavioral syndromes shape evolutionary trajectories via conserved genetic architecture

**DOI:** 10.1101/619411

**Authors:** Raphael Royauté, Ann Hedrick, Ned A. Dochtermann

## Abstract

Behaviors are often correlated within broader syndromes, creating the potential for evolution in one behavior to drive evolutionary changes in other behaviors. Despite demonstrations that behavioral syndromes are common across taxa, whether this potential for evolutionary effects is realized has not yet been demonstrated. Here we show that populations of field crickets (*Gryllus integer*) exhibit a genetically conserved behavioral syndrome structure despite differences in average behaviors. We found that the distribution of genetic variation and genetic covariance among behavioral traits was consistent with genes and cellular mechanisms underpinning behavioral syndromes rather than correlated selection. Moreover, divergence among populations’ average behaviors was constrained by the genetically conserved behavioral syndrome. Our results demonstrate that a conserved genetic architecture linking behaviors has shaped the evolutionary trajectories of populations in disparate environments—illustrating an important way by which behavioral syndromes result in shared evolutionary fates.

## Introduction

Behavior is frequently assumed to have been shaped by selection (Grafen 1984) and thus populations are expected to differ in a range of behaviors based on local selective pressures. This implies that behaviors are able to evolve independently, an assumption increasingly challenged by the ubiquity of behavioral syndromes—correlations among behaviors (Table 1; Sih et al. 2004a, Sih et al. 2004b)—which have been documented across taxonomic groups (Garamszegi et al. 2012, 2013) and are comprised of both genetic and environmental contributions (Dochtermann 2011, Dingemanse and Dochtermann 2014, Dochtermann et al. 2015).

**Table 1.**
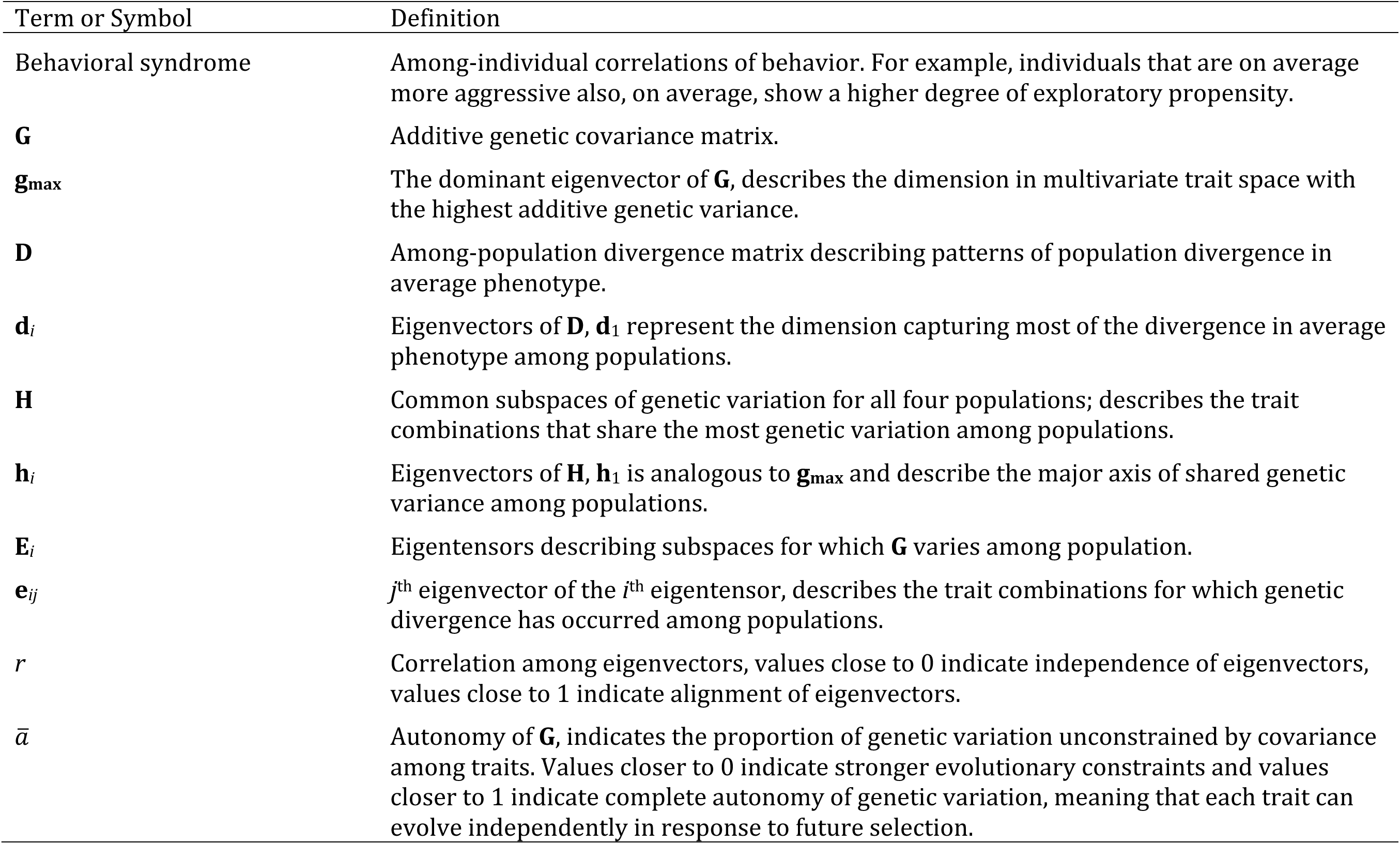
Terms and symbol definitions.

Given the contribution of genetic correlations to behavioral syndromes, these syndromes have the potential to constrain the ability of populations to diverge and respond to local selective pressures (Dochtermann and Dingemanse 2013). Specifically, based on quantitative genetic theory, if syndromes stem from pleiotropic effects—wherein a single gene affects multiple behaviors—populations will be constrained to diverge along shared evolutionary pathways. Further, this divergence is predicted to occur in the direction in trait space that contains the most variation (Figure 1; Schluter 1996). As a result, if syndromes have a constraining effect on evolution, the pattern of correlations among traits will be conserved among populations (Figure 1). Alternatively, if genetic correlations underpinning syndromes are the result of selection historically favoring particular trait combinations (i.e. selection-induced linkage disequilibrium (Roff 1997, Saltz et al. 2017)), the divergence of populations will be generally unconstrained as these genetic correlations are expected to rapidly break down when selection changes (Roff 1997, Conner 2002, Saltz et al. 2017).

**Figure 1.**
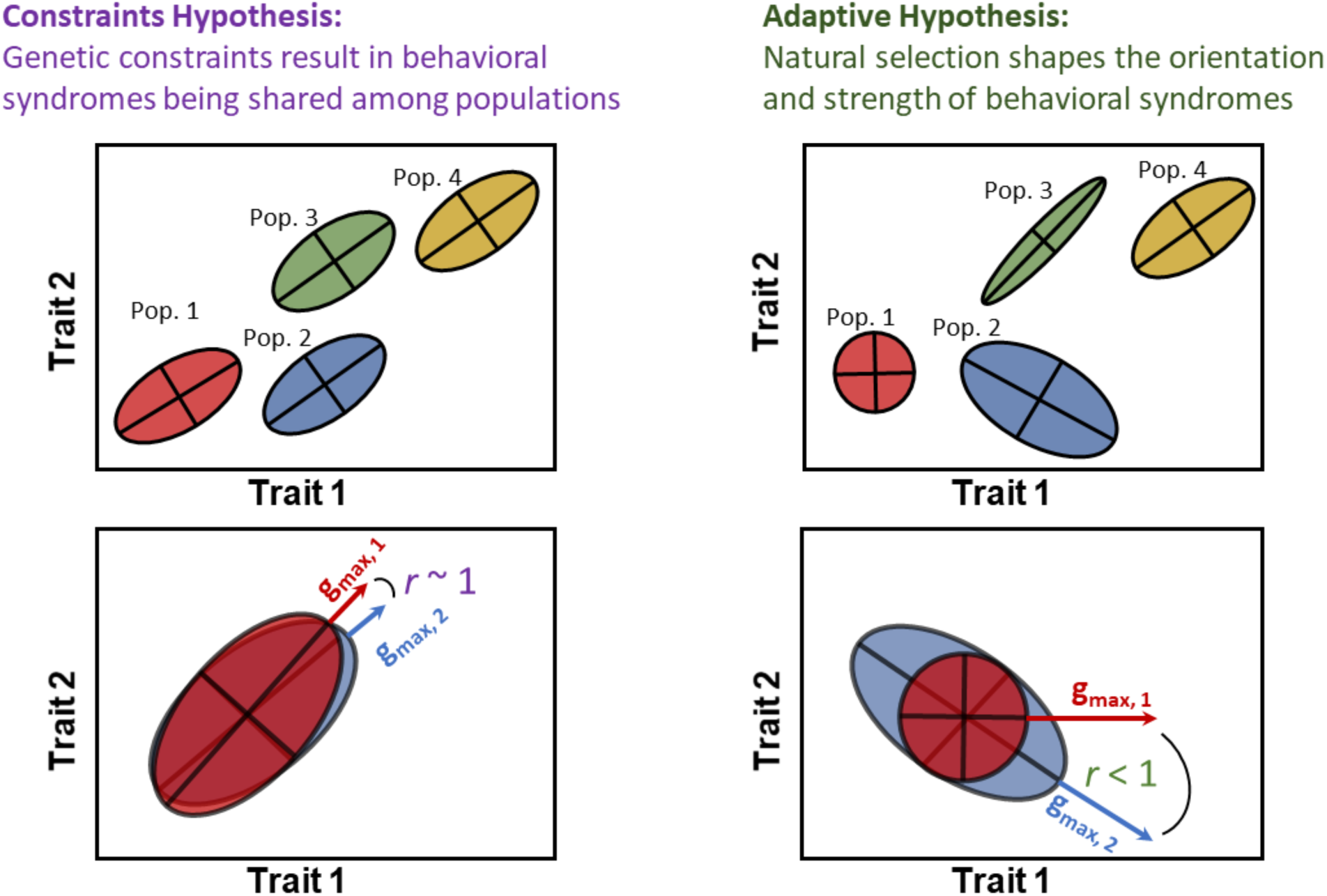
Two contrasting hypotheses can explain the presence of genetic correlations among behavioral traits (i.e. behavioral syndromes): Genetic constraints arising from pleiotropy and shared molecular mechanisms should lead to the expression of the same behavioral syndrome (top left panel). As a result, the vector correlations between major axes of genetic variation (**g_max_**) are predicted to be approaching 1 (bottom left panel). Alternatively, selection-induced linkage disequilibrium should lead to differing orientation and strength of behavioral syndromes when selective pressures differ among populations (top right panel). The vector correlation between **g_max_**s should therefore be below 1 (bottom right panel).

These two quantitative genetic explanations for behavioral syndromes have explicit analogs in the behavioral literature: whether syndromes emerge from pleiotropy, tight genetic linkage, or other shared physiological and cellular effects has been termed the “constraints hypothesis,” as opposed to selection-induced linkage disequilibrium, which has been termed the “adaptive hypothesis” (Figure 1, *sensu* Bell (2005)). While some studies have compared phenotypic or among-individual correlations across populations (e.g. Dingemanse et al. 2007, Pruitt et al. 2010, Dochtermann et al. 2012, Royauté et al. 2014, Michelangeli et al. 2018), only population comparisons of behavioral syndromes at the additive genetic level allow for properly testing these competing hypotheses. Unfortunately, data at the additive genetic level has been restricted to a single comparison of two populations (Bell 2005). Consequently, the overall role of behavioral syndromes in shaping population divergence is an important gap in our knowledge as support for the constraints versus adaptive hypotheses remains insufficiently tested.

Knowing whether behavioral syndromes emerge from genetic constraints or selection induced linkage disequilibrium is important because these mechanisms differentially affect evolutionary outcomes (Saltz et al. 2017). These potential effects are broad, from altering responses to environmental changes to speciation dynamics. For example, in *Anolis* lizards, constraints imposed by genetic correlations on morphological traits have shaped divergence and what phenotypes can be expressed during adaptive radiations (McGlothlin et al. 2018). Behavioral syndromes may have an even greater constraining effect: Dochtermann and Dingemanse (2013) reported that the average magnitude of genetic correlations between behaviors was sufficient to constrain evolutionary responses to a greater degree than do correlations between life-history or morphological traits. Unfortunately, this conclusion was based on data that could not distinguish between the constraints and adaptive hypotheses. If indeed the constraints hypothesis underpins behavioral syndromes then syndromes will reduce the rate of adaptation and reduce the rate at which populations and species diverge. Evaluating evidence for the constraints and adaptive hypotheses is therefore necessary in order to understand whether behavioral syndromes are important in behavioral evolution.

Here we evaluated predictions of the adaptive and constraints hypotheses (Figure 1) and tested whether behavioral syndromes have diverged at the genetic level among populations of the field cricket, *Gryllus integer*. Specifically, according to the constraints hypothesis we predicted that genetic variation would be expressed in a consistent manner among populations and genetic correlations would be maintained across generations. Specific predictions are more difficult to make for the adaptive hypothesis without knowledge of local selective pressures. This hypothesis can, however, be assessed indirectly because behavioral divergence is expected to not be constrained and correlations are expected to rapidly degrade. We evaluated these predictions via estimation and comparisons of behavioral genetic (co)variance matrices, i.e. **G** (Table 1), estimated for multiple populations of *G. integer* (Figure 2).

**Figure 2.**
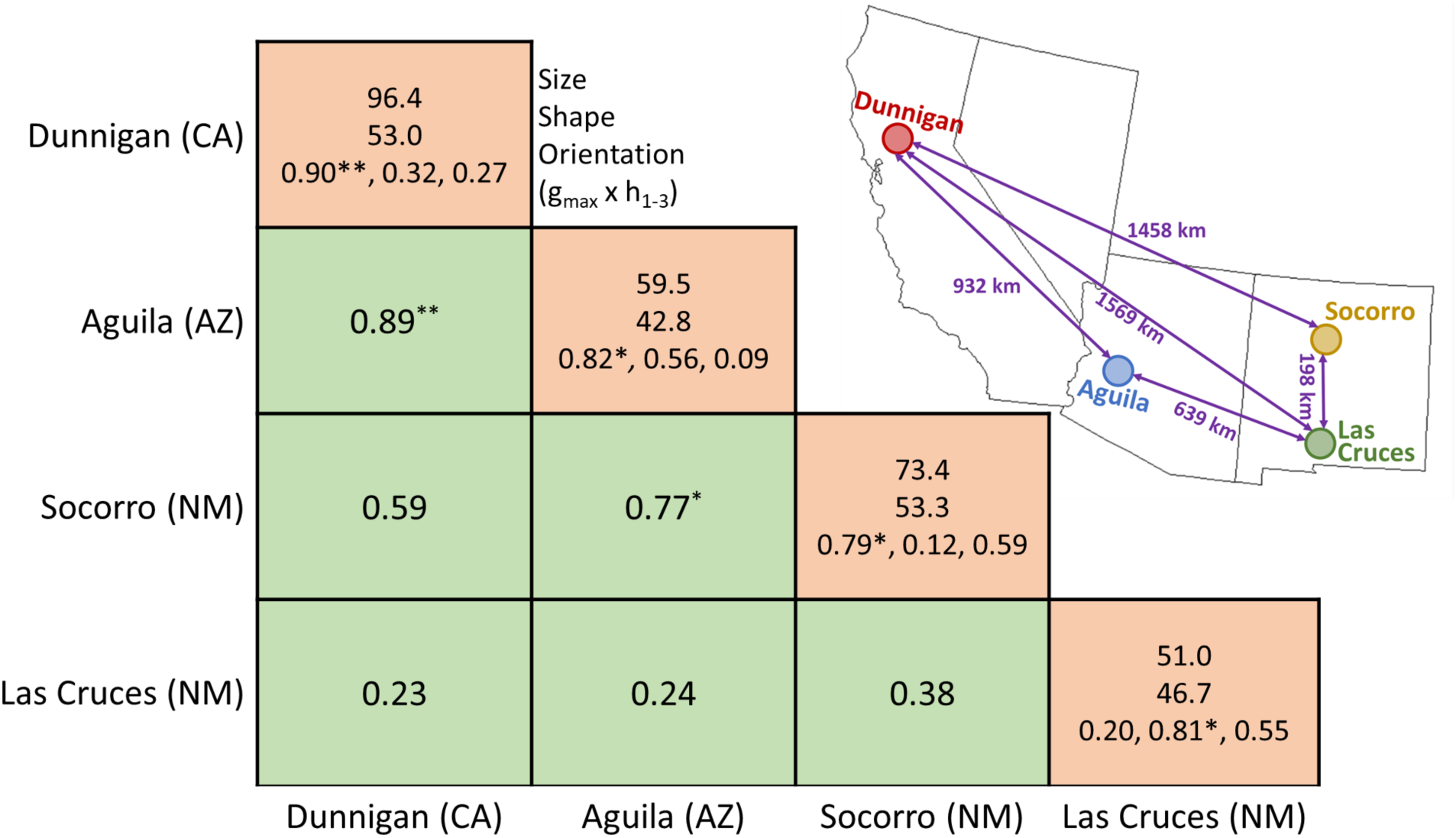
The genetic architecture of behavioral syndromes is shared in four isolated populations of *Gryllus integer*. Values along the diagonal (peach shading) describe the multivariate structure of behavioral variation: first row the size (total genetic variance), second the shape (percent of variance explained by the major axis of genetic variation, **g_max_**), and, third, orientation (vector correlation between **g_max_** and conserved genetic subspaces **h**_1-3_). Off-diagonal elements represent the correlation between the **g_max_** of each population (top row) and the probability that alignment differed from 0: ** P < 0.01, * P < 0.05.

## Methods

### Cricket collection

*G. integer* is a particularly appropriate species to evaluate the constraints and adaptive hypotheses for behavioral syndromes as the species exhibits population differences in ecologically relevant behaviors that can be assayed in the laboratory (e.g. Hedrick and Kortet 2006, Kortet and Hedrick 2007, Niemela et al. 2012a). *G. integer* can also readily be bred in the lab according to quantitative genetic designs making it an ideal model for our questions of interest (Hedrick 1988).

We collected adult female crickets from four populations throughout the southwestern and western US: Socorro, NM; Las Cruces, NM; Aguila, AZ; and Dunnigan, CA (Figure 2) during the summer of 2017. Crickets from these locations are formally recognized as members of *Gryllus integer* but additional splitting is currently being considered (D. Weissman, personal communication). These populations are also geographically distant from each other (Figure 2) and vary in predator and parasitoid abundance (Hedrick and Kortet 2006).

Around 50 females on average were collected from each population (Table S1) and taken to animal housing facilities at North Dakota State University. Females were housed individually in 0.71 L containers and provided with ad libitum food (Purina Chick Starter) and water (water was provided in glass vials capped with cotton). Each cricket was also provided with a small piece of cardboard egg carton for shelter. The cricket housing room was maintained on a 12:12 dark:light cycle reversed such that the room was dark during daytime hours. The housing room was kept at ∼27C.

### Breeding design

Females collected from the field (generation P) were allowed to oviposit in water vials while in their containers. Offspring of these females were designated generation F_0_ as sires were unknown: mating occurred prior to capture and multiple mating is common in the genus (Simmons 1986). F_0_ offspring hatched in their dams’ containers and were then moved to individual housing prior to maturation. We assayed the behavior of 387 F_0_ individuals (see below) upon maturation (Table S1). After behavioral trials, F_0_ individuals were assigned to breeding pairs such that individual males were mated to multiple randomly assigned females from the same population but different dams according to a standard full-sib, half-sib breeding design (Lynch and Walsh 1998). Matings were conducted as follows: females were moved from their normal housing containers to a larger container (34.6 × 21 × 12.4 cm) along with their food dish, water vial, and egg carton shelter. After the female had been transferred, the assigned male was likewise moved to the large container, also with its food dish, water vial, and egg carton. The male and female remained in these containers for 24 hours to allow sufficient time for courtship and multiple mating. After 24 hours the male and female crickets were returned to their original containers. If males were to be mated with additional females, they were allowed a minimum break of 24 hours before repeating the above procedure. These F_0_ females were subsequently allowed to oviposit into water vials within their containers. Resulting F_1_ offspring were moved to individual housing prior to maturation and had their behaviors assayed upon maturation. After behavioral assays, F_1_ individuals were likewise paired with F_1_ individuals of the same population but different sires in the same manner as above and resulting F_2_ offspring were moved to individual housing and had their behavior measured upon maturation. This resulted in the behavioral testing of 395 F_1_ individuals and 163 F_2_ individuals (Table S1). Across the three generations this represented behavioral testing of 946 individual crickets.

### Behavioral testing

All behavioral tests followed standard procedures previously validated in the literature for Gryllid crickets (Kortet and Hedrick 2007, Kortet et al. 2007, Hedrick and Kortet 2012, Niemela et al. 2012b, Royauté et al. 2015, Royauté and Dochtermann 2017, Royauté et al. 2019). These assays encompass how individuals vary in risk-taking behavior (Kortet et al. 2007), exploratory propensity (Royauté et al. 2015, Royauté and Dochtermann 2017, Royauté et al. 2019), and response to predation threat (Royauté and Dochtermann 2017, Royauté et al. 2019). Based on the known relatedness among individuals we then estimated **G**—the matrix of additive genetic behavioral variances and covariances—for each population. Below, we describe these behavioral assays and their ecological relevance.

#### Latency to emerge from shelter

Gryllid crickets, including *G. integer*, use small burrows and natural cracks for refuge from predators and to which they retreat when under threat. As a result, latency to emerge from shelter after disturbance can be considered a proxy for risk-taking behavior or “boldness” (Kortet et al. 2007). Here, we conducted latency tests wherein individuals were transferred from their home containers to small artificial burrows (40 cm^3^) placed within a 34.6 × 21 cm arena. These artificial burrows were capped so that individuals could not immediately emerge. Crickets were forced to remain in the artificial burrow for two minutes after which the cap was removed. Crickets were then allowed six minutes and thirty seconds to emerge from the artificial burrow. During this test we recorded how long it took for an individual to emerge (in seconds). Individuals that did not emerge were given a maximum latency of 390 seconds.

#### Open field exploratory behavior

Open field tests are a classic behavioral assay across taxa (Walsh and Cummins 1976) which measure the exploratory propensity of individuals (Réale et al. 2007), including crickets (Royauté et al. 2015, Royauté and Dochtermann 2017, Royauté et al. 2019). Individuals that move through more of the arena are considered more thorough explorers (Réale et al. 2007). Here we used open field tests to measure activity and exploratory propensity in a 30 × 30 cm plexiglass arena. Individuals were introduced into the arena under a small container and allowed to rest for 30 seconds after introduction. At the end of this 30 seconds, the container was removed and the cricket was allowed to explore the arena for 3 minutes and 40 seconds. The arena was cleaned with isopropyl alcohol between trials to remove any chemosensory cues from the arena. We used Ethovision XT to record the total distance the individual moved during the trial (cm), the number of unique zones of the arena an individual visited during the trial, and the variance in velocity of individuals. This latter measure indicates whether an individual’s speed of exploration was constant (low velocity variance) or whether individuals had frequent activity bursts punctuated by long bouts of inactivity (high velocity variance).

#### Response to cues of predator presence

How individuals respond to cues of predator presence often varies within and among populations and is likely to covary with fitness (Herman and Valone 2000). Crickets respond to chemical cues of predator presence by either freezing or increasing activity depending on whether confronted by predator cues of sit-and-wait or active predators (Storm and Lima 2008, Binz et al. 2014). Here we used a behavioral assay to measure response to cues of predator presence also previously used with another Gryllid species (Royauté and Dochtermann 2017, Royauté et al. 2019). Specifically, individuals were introduced into a 15 cm diameter circular arena (7.5 cm height), the floor of which was covered with dry filter paper that had been soaked with diluted excreta from leopard geckos (*Eublepharis macularius*). Crickets respond to exposure to leopard gecko cues by increasing activity (Royauté and Dochtermann 2017, Royauté et al. 2019). All leopard geckos were fed a diet of *G. integer* with occasional diet supplementation of mealworms (i.e. larval *Tenebrio molitor*) and the related decorated cricket (*Gryllodes sigillatus*). Crickets were introduced to a portion of the arena without predator cue under a small shelter. After a 30 second rest period, the shelter was removed and the individual allowed to freely move throughout the arena for 3 minutes and 40 seconds. We then used Ethovision XT to record the total distance an individual moved during the trial (cm). Total distance moved during the predator cue trial, the latency to first movement (in seconds), and the variance in velocity were used in subsequent analyses.

### Statistical analyses

#### **G** matrix estimation

We used multi-response mixed effect animal models (Kruuk 2004) implemented using the `MCMCglmm` package in R (Hadfield 2010) to estimate genetic variances and covariances (i.e. the **G** matrix). We included the effects of temperature, day and time of testing in the behavioral arena room along with sex, life-stage and mass of the individual as fixed effects. We used the individual relatedness matrix (based on the known pedigree) as a random effect. Traits for which variances and covariances were estimated were: (i) the latency that an individual emerged from the shelter during the trial (modeled as censored Gaussian), (ii) the distance moved during the open field trial (Gaussian), (iii) the number of unique zones an individual visited during the open field trial (Poisson), (iv) the log-transformed variance in velocity during the open field trial (Gaussian), (v) the square-root transformed distance an individual moved during the predator cue response trial (Gaussian), (vi) the latency to initiate movement in the antipredator response trial (Poisson) and (vii) the log-transformed variance in velocity during the antipredator response trial (Gaussian). The inclusion of dam ID as a random effect did not improve model fit, indicating negligible or non-existent maternal effects and was not included in final model runs. Multi-response models were fit individually by population with each population’s variances and covariances estimated from the posterior of an MCMC chain of 4.8 × 10^6^ iterations, with an 800,000 burn-in period and a thinning interval of 4,000. A prior that was minimally informative for both variances and covariances was used. All variances and covariances were estimated at the additive genetic level and on the latent scale (Table S4).

#### Evaluating the constraints and adaptive hypotheses

The location, orientation, and distribution of genetic variation in multivariate space—in our case seven dimensional space—can differ among groups in a variety of complex ways (Phillips and Arnold 1999, Blows et al. 2004, Walsh 2007, Roff et al. 2012, Aguirre et al. 2014). We therefore used a suite of statistical approaches to compare the orientation of genetic variation and evaluated agreement among multiple statistical approaches in support for either the constraints or adaptive hypothesis.

To determine whether behavioral syndrome structure at the additive genetic level was shared among populations we used two approaches:

i. testing whether populations exhibited shared subspaces (dimensions) of **G** based on Krzanowski’s common subspace analysis (Aguirre et al. 2014);
ii. comparing alignment of dominant eigenvectors among populations (i.e. **g_max_**, Table 1 (Schluter 1996))

Krzanowski’s common subspace analysis determines whether genetic variation is expressed in the same dimensions and direction across groups (Aguirre et al. 2014). This can be thought of as analogous to asking whether the populations shared principal components (Phillips and Arnold 1999). In two dimensions, this is similar to the directions the ellipses point in Figure 1, with shared subspaces when the ellipses point in the same direction. For our data there were seven possible dimensions of overlap but some dimensions may not possess substantive variation (Table S4). Following Aguirre et al. (2014) we therefore considered those subspaces that additively contained greater than 90% of the genetic variation, which were then summarized in matrix form (**H**)(Aguirre et al. 2014). Here, this included three possible shared subspaces (**h_1_, h_2_,** and **h_3_**; Table 1) and we tested whether these subspaces were shared among populations to a degree greater than expected by chance. Under the constraints hypothesis we expect genetic variation to be expressed in the same dimensional direction among populations, manifested as shared genetic subspaces. A lack of shared subspaces would therefore contradict the predictions of the constraints hypothesis. However, shared subspaces may also be observed under the adaptive hypothesis if selective pressures are the same across populations.

Because evolutionary trajectories are biased by the dimensional direction (vector) in which most genetic variation is expressed, we also compared this vector, i.e. **g_max_** (Schluter 1996), among populations. This approach is similar to Krzanowski’s common subspace analysis but focuses on the dimension in which most variation is expressed. Here we calculated the vector correlation between the **g_max_**s of each population and, via randomization testing, determined whether these correlations significantly differed from 0.

Even if genetic covariances constrain the path of evolutionary change, average behaviors can change. Therefore, we estimated “**D**” (Table 1)—a matrix that describes the phenotypic divergence amongst populations and was here estimated as the (co)variance of species means (Schluter 1996, McGlothlin et al. 2018). The eigenvectors (**d**, Table 1) of this matrix describe the direction in multivariate space in which most divergence has occurred. If behavioral syndromes have constrained the evolution of populations, we would expect the direction in which most phenotypic divergence in average behavior has occurred (**d_1_**) to be correlated with shared subspaces identified by Krzanowski’s common subspace analysis (McGlothlin et al. 2018). Here we calculated the vector correlation between **d_1_** and **h_1:3._** Non-zero correlations between **d_1_** and any shared subspace would be consistent with the constraints hypothesis.

As with average behaviors, genetic variances across dimensions may differ among populations even while populations share an overall conserved genetic behavioral syndrome. We can therefore ask whether genetic variances have been constrained by a conserved behavioral syndrome. To do so we used methods described by Hine et al. (2009) to calculate what are known as genetic covariance eigentensors (**E**, Table 1). This approach starts by calculating the variances and covariances of **G** matrices across populations. Resulting matrices can subsequently be subjected to further eigen analysis, producing eigentensor matrices (**E**) and eigenvectors (**e**, Table 1) to describe in which dimension the **G**s of populations differ the most. If a conserved genetic behavioral syndrome, consistent with the constraints hypothesis, has affected the divergence of **G**s among populations we would expect correlations between the eigenvectors of eigentensors (**e**) and any shared subspaces (**h_1:3_**). Put another way, if populations share genetic variation in the same dimension in which most of the genetic divergence occurred, this would be evidence that syndromes channel behavioral evolution along a line of least resistance.

For the above approaches we followed the recommendations of Aguirre et al. (2014) in that all tests were based on the full MCMC posterior distributions and null distributions for population comparisons were based on randomizations of breeding values. To compare whether eigenvectors were significantly aligned, we also generated a random distribution of vector correlations following McGlothlin et al. (2018). The critical values of vector correlations based on this distribution were 0.93 (P < 0.001), 0.85 (P < 0.01), 0.71 (P < 0.05) and 0.62 (P < 0.1). To assess the significance of eigenvalues of **H** and **E** against random expectations, we calculated the largest posterior quantiles for which these distributions did not overlap (Figures S2 and S3 respectively). This threshold serves as a Bayesian probability in favor of the observed distribution being generated by patterns other than chance (hereafter, P_mcmc_). Because there are no clear-cut rules for interpreting these Bayesian probabilities, we provide the following scale to indicate how we interpreted support for inferences: P_mcmc_ < 0.7: poor evidence of difference compared to random expectations; P_mcmc_ > 0.8: moderate evidence of difference compared to random expectations; P_mcmc_ > 0.9 strong evidence of difference compared to random expectations; P_mcmc_ > 0.95: very strong evidence of difference compared to random expectations. Other reported probabilities (hereafter P) were interpreted according to standard criteria.

To further assess support for the adaptive and constraints hypotheses we also compared genetic correlations across generations. Genetic correlations due to selection-induced linkage disequilibrium are expected to decline across generations with random mating. Specifically, Conner (2002) argued that with random mating and in the absence of physical linkage, the magnitude of genetic correlations should halve every generation. Because of our breeding design we were able to opportunistically and separately estimate phenotypic and genetic correlations among behaviors by generation. Here, while mating was restricted to be within populations, mating was random with regard to behavior and we would therefore expect both genetic correlations (*r*_A_) and phenotypic correlations (*r*_P_) to decrease during the duration of the experiment under the adaptive hypothesis. We therefore generated expected genetic and phenotypic average absolute correlations under these assumptions (Appendix S1) and compared the observed estimates to these expectations. According to the constraints hypothesis we would expect correlations to remain stable across generations while under the adaptive hypothesis we would expect them to degrade toward a correlation of zero.

Finally, based on the estimated **G** matrices for each population, we calculated “autonomy” (*ā*, Table 1) throughout multivariate space following Hansen and Houle (2008). Autonomy provides an estimate of the “fraction of genetic variation that is independent of potentially constraining characters”(Hansen and Houle 2008). Put another way, autonomy estimates the degree to which genetic variation is free to respond to selection (max *ā*= 1) versus constrained by covariance (min *ā*= 0). We did not have predictions as to values for autonomy under either the constraints or adaptive hypotheses. Instead, these values indicate the potential for future evolutionary constraints for each population.

## Results

Behavioral syndromes were genetically conserved among populations. Based on Krzanowski’s common subspace analysis (**H**, Table 1; Aguirre et al. (2014)), the behavioral syndrome of *G. integer* was characterized by three dimensions of genetic covariance (**h**_1-3_, Table 1). These dimensions, and thus the overall syndrome, were shared among populations, as indicated by all Bayesian probabilities, p_mcmc_, being < 0.65 (Fig. S1). p_mcmc_ values that are closer to 1 for this test would indicate departure from random expectations and would therefore support a lack of shared syndrome structure among populations. The shared behavioral syndrome was comprised of: *i*) genetic covariation between shelter emergence time and predator cue responsiveness (**h**_1_, Table 2); *ii*) a genetic boldness-activity syndrome in which active individuals were more prone to ignore predator cues and were quicker to exit from their shelter (**h**_2_, Table 2); and *iii*) genetic covariance between activity and shelter emergence (**h**_3_, Table 2). Each of these three axes explained around one-third of the observed genetic variance (Table 2).

**Table 2.** Eigenvectors of phenotypic divergence (**d**), conserved genetic variation (**h**) and divergence in **G** (**e**). Traits legend: Latency = latency to exit from the shelter, OF.Dist = distance travelled in the open-field test, UZ = number of unique zones explore in the open-field arena, OF.Var.Velo = variance in velocity in the open-field test, AP.Dist = distance travelled in the antipredator response test, AP.Lat.Mov = latency to initiate movement in the antipredator response test, AP.Var.Velo = variance in velocity in the antipredator response test.

| Traits | **d**<sub>1</sub> | **d**<sub>2</sub> | **h**<sub>1</sub> | **h**<sub>2</sub> | **h**<sub>3</sub> | E1 (53 %) | E2 (31 %) |  |
| --- | --- | --- | --- | --- | --- | --- | --- | --- |
|  |  |  |  |  |  | **e**<sub>11</sub> | **e**<sub>21</sub> | **e**<sub>22</sub> |
| Latency | **-0.32** | **0.90** | **-0.31** | **0.86** | **0.47** | **0.63** | 0.08 | **0.77** |
| OF.Distance | **-0.80** | **-0.34** | 0.18 | **-0.37** | **0.88** | -0.13 | **-0.79** | **0.42** |
| UZ | -0.02 | -0.06 | 0.02 | -0.02 | 0.06 | -0.01 | -0.08 | 0.06 |
| OF.Var.Velo | -0.09 | -0.06 | 0.02 | -0.02 | 0.07 | -0.02 | -0.10 | 0.06 |
| AP.Distance | **-0.48** | -0.04 | **0.93** | **0.35** | 0.02 | **-0.74** | **-0.59** | **-0.45** |
| AP.Lat.Mov | -0.05 | 0.24 | -0.10 | -0.01 | 0.02 | 0.16 | 0.07 | 0.15 |
| AP.Var.Velo | -0.07 | -0.03 | 0.05 | 0.02 | 0.01 | -0.05 | -0.05 | -0.02 |
| % Variance explained | 58.2 | 31.4 | 33.1 | 33.0 | 32.9 | 97.4 | 69.5 | 30.4 |

Following the demonstration of genetic conservation of behavioral syndromes, we determined whether genetic variation was primarily expressed in the same direction in multivariate space across populations (**g_max_** alignment; Table 1). Put another way, given the general conservation of behavioral syndrome structure at the genetic level, did populations express most genetic variation in the same combinations of traits? Indeed, the **g_max_**s of the Aguila and Dunnigan and Socorro and Dunnigan populations were strongly correlated with each other (vector correlation *r* > 0.7, p < 0.05) (Figure 2). Moreover, the **g_max_**s of each population were aligned with the shared axes (Figure 2). This alignment demonstrates that the genetically conserved behavioral syndrome captured the genetic variation expressed in each population and confirmed that the orientation of genetic variation in multivariate space was conserved among the populations.

Despite the genetic conservation of behavioral syndrome structure, populations did exhibit some divergence. Specifically, the populations have diverged in their multivariate behavioral averages (i.e. “**D,**” Figure 3, Table 1 and S2) and in the magnitude of genetic variation present in each population (Figure S3). Importantly, however, the direction of divergence in both means and variances was aligned with the shared behavioral syndrome (e.g. *r*_d1,h3_ = 0.85, p < 0.01; *r*_h1,e11_ = 0.92, p < 0.01; Table S2). This alignment demonstrates that divergence has been constrained by the shared structure of behavioral syndromes.

**Figure 3.**
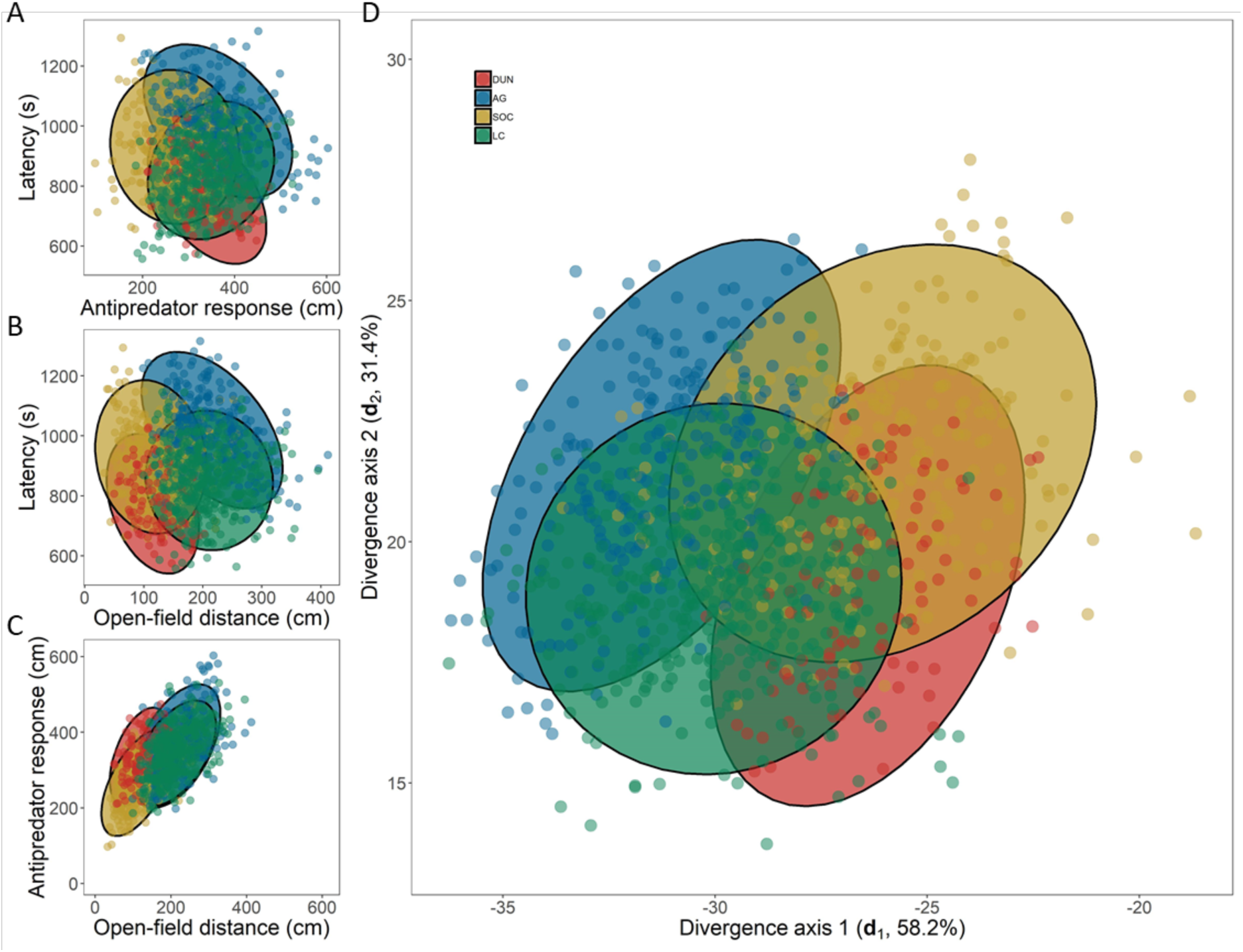
Evolutionary divergence in the structure of behavioral syndromes occurs along shared axes of genetic variation. A-C) Correlations between pairs of traits that exhibit the greatest variation in divergence (Table S2). Points represent breeding values for each individual within a population centered around the population mean for that trait. >50% of divergence was in latency to emerge from shelter by antipredator response activity D) Population-specific divergence in average behaviors. Population-specific **G** matrices were visualized by transforming estimated breeding values for each trait based on the divergence among populations. Ellipses represent the 95% confidence ellipses for each population centered at the multivariate species mean (DUN: Dunnigan CA, AG: Aguila AZ, SOC: Socorro NM, LC: Las Cruces NM).

Behavioral syndromes emerging from either the adaptive or constraints hypotheses are expected to respond differently to random mating. Specifically, under the adaptive hypothesis, genetic correlations are expected to erode by 50% every generation. Because we mated individuals at random, we were able to compare the observed average genetic and phenotypic correlations (*r*_A_ and *r*_P_) with their expected values under the adaptive hypothesis (see Appendix S1 for details). Contrary to the expectations of the adaptive hypothesis, but as predicted according to the constraints hypothesis, average genetic and phenotypic correlations remained stable over the course of three successive laboratory generations (posterior mean and 95 % credible intervals; *r*_A Observed_ F_1_ = 0.36 [0.23; 0.52], *r*_A Observed_ F_2_ = 0.38 [0.23; 0.53], Figure 4).

**Figure 4.**
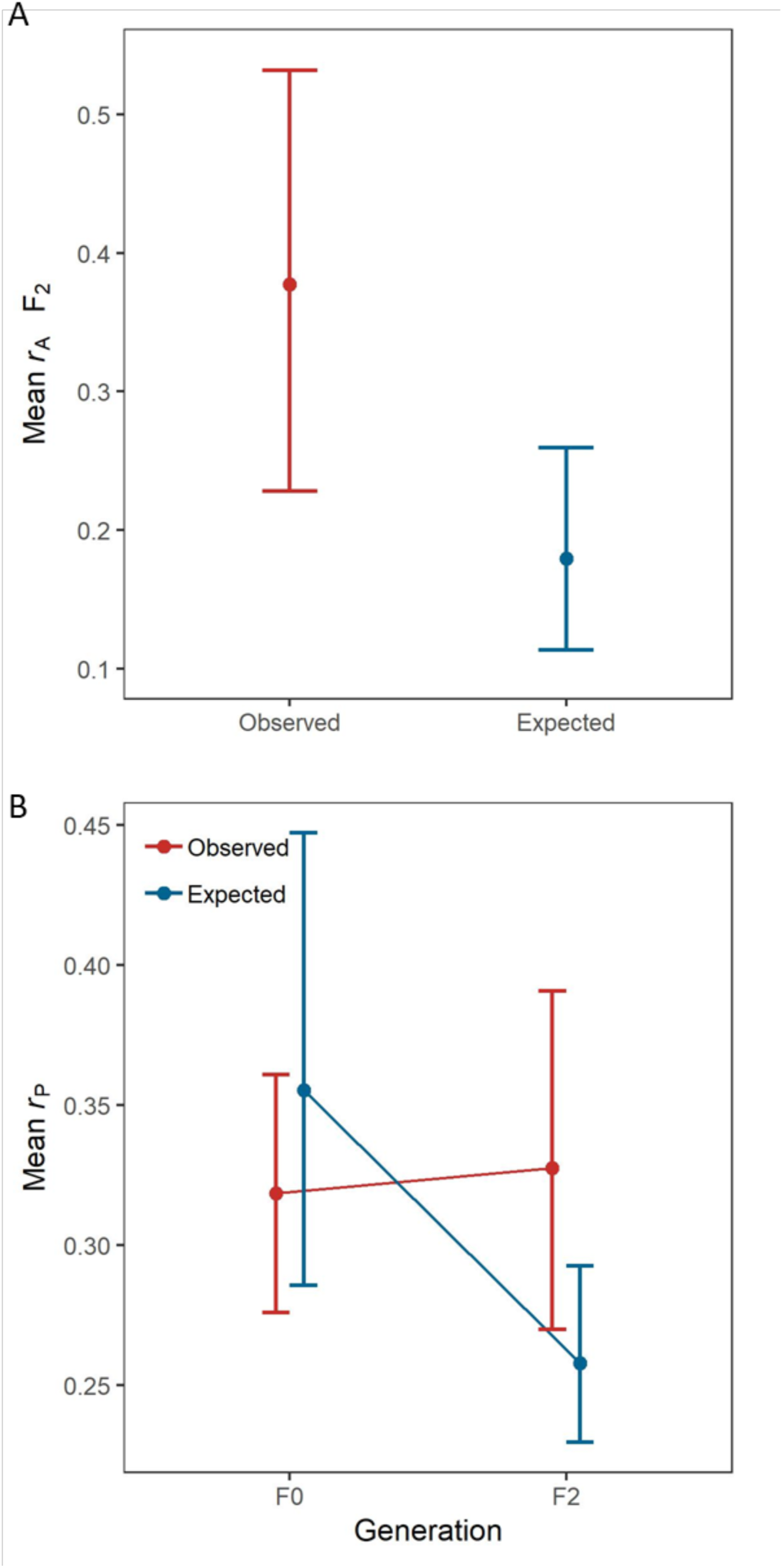
A) Additive genetic (*r*_A_) and B) phenotypic correlations (*r*_P_) remained stable over the course of three successive generations compared to theoretical expectations based on selection-induced linkage disequilibrium and random mating (*r*_A *observed*_ = 0.38, *r*_A *expected*_ = 0.18, P_mcmc_ for difference from expectations under selection-induced linkage disequilibrium > 0.85; *r*_P *observed*_ F_0_ = 0.32, *r*_P *observed*_ F_2_ = 0.33, *r*_P *expected*_ F_0_ = 0.35, *r*_P *expected*_ F_2_ = 0.26, P_mcmc_ F_2_ > 0.80). Error bars correspond to 95% credibility intervals around the posterior mean.

Finally, for each population, we calculated autonomy (Table 1), which estimates the degree of constraint on evolutionary outcomes imposed by the genetic architecture connecting traits. Autonomy varies between 0 and 1, with higher values indicating greater potential for independent evolution. For *Gryllus integer*, autonomy varied between 0.47 and 0.61 (DUN: *ā*= 0.48 [0.31; 0.68], SOC: *ā*= 0.47 [0.32; 0.67], AG: *ā*= 0.60 [0.44; 0.78], LC: *ā*= 0.61 [0.43; 0.76], all populations combined: *ā* = 0.57 [0.43; 0.70]). This suggests that the constraining effect of behavioral syndromes is likely to persist over future generations.

## Discussion

Three key results demonstrate conservation of behavioral syndromes at the genetic level despite differences among populations in average behavior, providing strong support for the constraints hypothesis. This support for the constraints hypothesis is unexpected given that a previous study with stickleback (Bell 2005) found that two populations differed in the magnitude of heritabilities and genetic correlations between two behaviors—albeit with overlapping confidence intervals—providing support in that case for the adaptive hypothesis. This conservation of behavioral syndrome structure has also had the effect of channeling population divergence. Our results therefore suggest that studying a broader suite of behavioral traits may reveal evolutionary constraints not apparent from pair-wise correlations.

Our first major result supporting the constraints hypothesis was that the genetic variation among the four populations was shared along three dimensions. These dimensions describe the genetic structure of the species’ behavioral syndromes and their being shared demonstrates that the orientation of genetic variation was conserved among populations. The overall behavioral syndrome consisted of a boldness-activity dimension (**h**_2_, Table 2) frequently described in the literature. This dimension genetically links activity, exploration and risk-prone behaviors. This dimension has been described at the phenotypic level (Wilson and Godin 2009, Bókony et al. 2012) but demonstrations at the genetic level are rare (see Bell (Bell 2005) for one example). The other conserved dimension (**h**_1_ and **h**_3_) represent potential trade-offs between risk management strategies, in which individuals either compensate for risk during foraging by being less prone to resume activity when threatened (**h**_3_, Table 2), or take risks in one context (not moving away from a predator cue) while avoiding risk in another (taking longer to emerge from shelter) (**h**_1_, Table 2). Alternatively, dimension **h**_1_ might indicate that individuals with long latencies are less active. As a result, these individuals may encounter fewer predator cues resulting in weaker antipredator responses.

Our second major result supporting the constraints hypothesis was that the **g**_max_s of the Aguila and Dunnigan and Socorro and Dunnigan populations were strongly correlated and all **g**_max_s were aligned with the shared behavioral syndrome. This validates that behavioral syndrome structure is shared among the populations and that the behavioral syndrome captures the majority of observed genetic variation. Schluter (1996) demonstrated that morphological divergence among several pairs of populations and species of vertebrates is constrained by **g**_max_. Specifically, evolutionary divergence was greatest when populations and species shared a common **g**_max_ and there was directional selection for morphological trait combinations in this same direction in phenotypic space (Schluter 1996). We found that **g**_max_ was conserved and that divergence in both average behavior and genetic (co)variance among the four populations was aligned with **g**_max._. This demonstrates that behavioral syndromes affect population divergence in a manner similar to that observed for morphology.

Our third result in support of the constraints hypothesis stems from the prediction that, under the adaptive hypothesis, genetic correlations are expected to decrease by about 50% each generation due to the effects of recombination (Conner 2002). This prediction assumes an absence of genetic linkage and random mating (Appendix S1). However, genetic linkage sufficiently strong to resist recombination is also consistent with the constraints hypothesis—see, for example, the effects of supergenes (Purcell et al. 2014, Küpper et al. 2016)—and so we consider this assumption appropriate. In contrast to this prediction of declining correlations, we found that the average genetic correlation did not change across generations (Figure 4). Similarly, phenotypic correlations did not decrease according to predictions (Figure 4). Because we were not able to study replicate lines under random mating, this finding is not conclusive on its own. Instead the result is one additional line of evidence consistent with the constraints hypothesis and in contradiction of the adaptive hypothesis.

Importantly, the first two results—shared dimensions of genetic variation and correlated **g**_max_s—could also be observed under the adaptive hypothesis if the selective pressures each of the populations experienced were the same. We consider this unlikely for three reasons. First, the degree of geographic separation among populations was extensive, totaling more than 1500 km in some cases (Figure 2). This degree of geographic separation makes it unlikely that the populations experienced the exact same selective regime. Moreover, climate (Table S5) as well as predation and parasitism regimes are highly variable among the populations (Hedrick and Kortet 2006). Second, if similarity in selection regimes was the driving force behind these converging patterns of genetic covariance, we would expect the geographically closest populations to have the greatest similarity in **g**_max_. This was not the case and, in fact, **g**_max_ was most similar among populations that were geographically most separated (Figure 2). Finally, our third main result directly contradicts the adaptive hypothesis: if trait correlations, like those of behavioral syndromes, arise due to the adaptive hypothesis and therefore selection-induced linkage disequilibrium, they are expected to rapidly degrade under random mating (Roff 1997, Conner 2002). In direct contradiction to this expectation we observed that correlations did not decrease across generations (Figure 4). Put another way, our first two results—which showed that the multivariate composition of behavioral syndromes was shared among populations—are consistent with the predictions of the constraints hypothesis. Next, our third result—the maintenance of behavioral correlations despite random mating—demonstrates the failure of predictions made by the adaptive hypothesis.

Our results indicate that the conserved genetic architecture of behavioral syndromes leads to populations having quantitatively constrained evolutionary trajectories (Houle 2001) and that these syndromes have limited population divergence. This quantitative constraint and resulting limitation on divergence is also expected to persist into the future due to the behavioral syndrome structure imposed by each population’s **G** matrix. Based on these **G** matrices, we found similar degrees of autonomy (Hansen and Houle 2008) among populations ranging from 0.47-0.61, a stronger constraint than observed for life-history or morphological traits (Dochtermann and Dingemanse 2013). These autonomies indicate that behaviors will rarely evolve independently and the observed genetic behavioral syndrome will affect future evolution.

Despite the conservation of behavioral syndrome structure at the genetic level across populations, *Gryllus integer* populations did exhibit divergence in both mean behaviors and magnitudes of genetic variation present in each population. Both the divergence in means and variances was strongly aligned with the shape of the shared behavioral syndrome, demonstrating that the syndrome itself was channeling the evolutionary divergence of the populations (Table S2). The divergence in means was most strongly related to differences in latency and distance moved in both the open-field and antipredator assays (Table 2). Specifically, the Dunnigan, CA and Aguila, AZ populations exhibited the greatest differences in average behaviors (Figure 4). Similarly, the divergence in magnitude of genetic variation was driven by the three easternmost populations having less genetic variation than the Dunnigan, CA population (Figure S1). Whether this represents a loss of variation due to selection, stochastic effects on the three eastern populations, or the accumulation of variation for the western population is not currently clear.

Throughout we have referred to the adaptive and constraints hypothesis as competing hypotheses for the expression of behavioral syndromes. However, two caveats to this framing exist: First, according to a Tinbergian framework (Tinbergen 1963), these hypotheses are not addressing questions at the same level. Within the framework of Tinbergen’s four questions, the constraints hypothesis is a proximate causation question and reduces to a question about pleiotropy (or other molecular mechanisms) versus linkage and is agnostic as to selection. In contrast, the adaptive hypothesis is an ultimate question of function and would be assessed by determining the alignment between **G** and selection gradients (e.g. following methods described by Berdal and Dochtermann 2019). This framing does not, however, carry-over to the quantitative genetic literature wherein selection-induced linkage disequilibrium and molecular mechanisms such as pleiotropy are considered competing explanations (e.g. Roff 1997). The difference in perspective is partly due to the history of the fields but is also because the quantitative genetic framework recognizes the necessary role of continuing correlated selection in maintaining covariances stemming from linkage disequilibrium. As a second caveat it is important to note that the hypotheses are not strictly mutually exclusive (Conner et al. 2011, Saltz et al. 2017). It is possible that some portion of an estimated genetic covariance might stem from pleiotropy while some other portion stems from linkage disequilibrium (the two could even cancel each other out). Nonetheless, our results consistently supported the predictions of the constraints hypothesis across several lines of evidence.

The surprising degree of shared genetic variation in behavioral syndrome reported here suggests an unrecognized and important role for behavioral syndromes in the evolution of populations. Behaviors such as those measured here—exploratory behaviors and responses to predation threat—are frequently assumed to have been under selection and their responses to selection have been assumed to be unconstrained. In contrast, we have shown that the genetic contribution to behavioral expression is highly conserved, that populations share evolutionary fates, and that conserved behavioral variation may be a driver of population divergence and perhaps even speciation.

## Acknowledgements

We thank Monica Berdal, Katelyn Cannon, Jeremy Dalos, Sarah Felde, Brady Klock, Ishan Joshi, Hannah Lambert, Jenna LaCoursiere and Alondra Neunsinger for assistance in conducting behavioral trials and in rearing and care of the crickets and Martori Farms, David Lightfoot, Scott Bundy, Nico Franz, Sangmi Lee, Cameron Jones, Kenny Chapin, Ti Eriksson, Meranda Feagins, Charlotte Mulloney, Melody Martinez, Allyson Richins, Mauriel Rodriguez, Helen Vessels and David Wikman for assistance in collecting the crickets. We thank J. Bowsher, J. Pruitt, D. Westneat, and members of the Gillam and Dochtermann labs at NDSU for important discussions and/or comments on earlier versions of this paper. We also thank Adam Reddiex and two anonymous reviewers for essential critical feedback. This work was supported by US NSF IOS grants 1557951 and 1558069 to N.A.D. and A.H. respectively.

## Notes

#### Summary of Updates

Revised version following reviewer comments

https://github.com/DochtermannLab/G-PopComparison

