## Supplementary appendix (S1) tables (S1-S5) and figures (S1-S3) for "Behavioral syndromes shape evolutionary trajectories via conserved genetic architecture"

Raphaël Royauté, Ann Hedrick, and Ned A. Dochtermann

**Appendix S1.** Comparing observed patterns of phenotypic correlations with expectations based on SILD and random mating.

**Table S1.** Sample sizes by population and generation.

**Table S2.** Vector correlation (*r*) between major axes of phenotypic divergence (**d**), subspaces of conserved genetic variation (**h**) and subspaces representing genetic divergence (**e**).

**Table S3.** Genetic variances and covariances of four populations of field crickets.

**Table S4.** Eigen decomposition of **G** for each population.

**Table S5**. Annual temperatures and precipitation

**Figure S1.** Population coordinates and genetic variance along the leading eigentensors capturing most of the variation in divergence (**E**1 and **E**2).

**Figure S2.** Eigenvalues of **H** for a comparison of the first three eigenvectors of each **G** matrix compared against random expectations.

**Figure S3.** Eigenvalues of the first three positive eigentensors of **S**.

**Appendix S1.** Comparing observed patterns of phenotypic correlations with expectations based on SILD and random mating.

Genetic correlations are expected to decline every new generation when they arise from selection-induced linkage disequilibrium (SILD). Conner [11] showed that in absence of both selection and linkage, the magnitude of genetic correlations under a SILD model and random mating should halve with every new generation. In contrast, genetic correlations caused by pleiotropic effects are expected to remain stable even after multiple generations of random mating. From this we expect the additive genetic correlation in the $F_2$ generation to be half as large as that of the $F_1$ generation:

$$(1)\ r_{\mathrm{A,\,F_2}} = \frac{1}{2} r_{\mathrm{A,\,F_1}}$$

To test this hypothesis, we estimated the observed genetic covariance based on multivariate animal models including the data of the $F_0$ and $F_1$ and $F_1$ and $F_2$ generations respectively. These models were specified similarly to the multivariate models described in the main text, but population origin was added as a fixed effect to control for population differences in average trait expression. We transformed the genetic covariance matrix to a correlation matrix and calculated the observed average correlation coefficients, $r_{\mathrm{A}\ observed}$, for every generation. We then compared, $r_{\mathrm{A}\ observed}$ to the expected average correlation shown in equation (1). Since our multivariate models were specified within a Bayesian framework, we performed these operations on every slice of the posterior estimates to obtain 95% credible intervals of observed and expected estimates and base our inference on the overlap between the two 95% credible intervals.

Because phenotypic correlations result from the combined influence of genetic and environmental influences, it is also possible to compare the expected decline in phenotypic correlations under SILD and random mating with the phenotypic correlations observed in every generation. The expected relationship between the phenotypic covariance in generation $F_0$, $F_1$ and $F_2$ can therefore be expressed by the following sets of equations:

$$(2)\ \mathbf{P}_{\mathbf{F_0}} = \mathbf{2}\ \mathbf{G}_{\mathbf{F_1}} + \mathbf{E}$$

$$(3)\ \mathbf{P}_{\mathbf{F_1}} = \mathbf{G}_{\mathbf{F_1}} + \mathbf{E}$$

$$(4)\ \mathbf{P}_{\mathbf{F_2}} = \frac{\mathbf{1}}{\mathbf{2}}\mathbf{G}_{\mathbf{F_1}} + \mathbf{E}$$

Where $\mathbf{P}_{\mathbf{F_x}}$, and $\mathbf{G}_{\mathbf{F_x}}$ represent the phenotypic and genetic covariance matrices for generation $F_x$ respectively and **E** represents the overall environmental covariance matrix estimated across all generations. Note that these formulas only apply to the covariance among traits (i.e. the off-diagonal elements) while the genetic variances (i.e. the diagonal elements) are kept constant across generations. $\mathbf{G}_{\mathbf{F_1}}$ was used as our reference here because this generation had the largest sample size and thus had the most influence on the estimation of **G**. We estimated $\mathbf{G}_{\mathbf{F_1}}$ from the animal model used to estimate equation (1) and we estimated **E** using a model containing the data from all populations for every generations.

We then compared these expected covariances to the observed phenotypic covariance estimated for each generation separately by fitting multivariate models excluding pedigree information so that only the phenotypic covariance was estimated. As above, we converted each covariance matrix to a correlation matrix and compared the observed average correlation coefficient $r_{P\ observed}$, with the average correlation coefficient $r_{P\ \mathrm{expected}}$ for every generation.

The full breakdown of these correlations is summarized below, see also Figure 3.

| Level | Generation | Observed Average $r$ | [95% CI] | Expected Average $r$ | [95% CI] |
|---|---|---|---|---|---|
| Genetic | | | | | |
| | $F_1$ | 0.36 | [0.23; 0.52] | - | - |
| | $F_2$ | 0.38 | [0.23; 0.53] | 0.18 | [0.11; 0.26] |
| Phenotypic | | | | | |
| | $F_0$ | 0.32 | [0.27; 0.36] | 0.35 | [0.28; 0.45] |
| | $F_1$ | 0.27 | [0.24; 0.31] | 0.29 | [0.25; 0.34] |
| | $F_2$ | 0.33 | [0.27; 0.39] | 0.26 | [0.23; 0.29] |

**Table S1.** Number of sires, dams and phenotyped offspring for each population and generation. The $F_0$ population represents the first population established in the laboratory after collection of gravid females in the field. N indicates the number field collected females for the $F_P$ generation.

| Population and generation | No. of sires | No. of dams | No. offspring in pedigree | No. phenotyped offspring |
|---|---|---|---|---|
| Dunnigan (CA) (N = 38) | | | | |
| $F_0$ | 0 | 8 | 30 | 30 |
| $F_1$ | 6 | 8 | 36 | 36 |
| $F_2$ | 6 | 8 | 35 | 35 |
| Total | 12 | 24 | 101 | 101 |
| Aguila (AZ) (N = 71) | | | | |
| $F_0$ | 0 | 32 | 144 | 143 |
| $F_1$ | 16 | 21 | 92 | 92 |
| $F_2$ | 2 | 2 | 16 | 16 |
| Total | 18 | 55 | 252 | 251 |
| Socorro (NM) (N = 65) | | | | |
| $F_0$ | 0 | 19 | 112 | 112 |
| $F_1$ | 10 | 20 | 126 | 125 |
| $F_2$ | 4 | 4 | 8 | 8 |
| Total | 14 | 43 | 246 | 245 |
| Las Cruces (NM) (N = 38) | | | | |
| $F_0$ | 0 | 13 | 105 | 102 |
| $F_1$ | 20 | 47 | 157 | 142 |
| $F_2$ | 13 | 16 | 104 | 104 |
| Total | 33 | 76 | 366 | 349 |

**Table S2.** Vector correlation (*r*) between major axes of phenotypic divergence ($\mathbf{d}_1$, $\mathbf{d}_2$), subspaces of conserved genetic variation ($\mathbf{h}_1$, $\mathbf{h}_2$, $\mathbf{h}_3$) and subspaces representing genetic divergence ($\mathbf{e}_{11}$, $\mathbf{e}_{21}$, $\mathbf{e}_{22}$). Probability of significant alignment: *** P < 0.001, ** P < 0.01, * P < 0.05. Italics indicate significant alignment at P < 0.1.

| | $\mathbf{d}_1$ | $\mathbf{d}_2$ | $\mathbf{h}_1$ | $\mathbf{h}_2$ | $\mathbf{h}_3$ |
|---|---|---|---|---|---|
| $\mathbf{h}_1$ | 0.49 | 0.40 | | | |
| $\mathbf{h}_2$ | 0.15 | 0.89** | | | |
| $\mathbf{h}_3$ | 0.85** | 0.06 | | | |
| $\mathbf{e}_{11}$ | 0.25 | *0.69* | 0.92** | 0.33 | 0.18 |
| $\mathbf{e}_{21}$ | 0.90** | 0.40 | 0.72* | 0.16 | *0.67* |
| $\mathbf{e}_{22}$ | 0.38 | 0.60 | 0.59 | 0.34 | 0.72* |

**Table S3.** Genetic variances (shaded diagonal elements) and covariances (bottom off-diagonal elements) of four populations of field crickets sampled in Dunnigan (CA), Aguila (AZ), Socorro (NM) and Las Cruces (NM). The probability of excluding zero (Pmcmc) is indicated on the top diagonal. Bold values indicate Pmcmc > 0.95, bold and italics indicate Pmcmc > 0.90 and italics indicate Pmcmc > 0.80.

| | Latency | OF.Dist | UZ | OF.Var.Velo | AP.Dist | AP.Lat.Mov | AP.Var.Velo |
|---|---|---|---|---|---|---|---|
| Dunnigan (CA) | | | | | | | |
| Latency | 33.66 | 0.58 | 0.66 | 0.61 | *0.88* | **0.95** | 0.77 |
| OF.Dist | 1.89 | 21.58 | **0.95** | **0.99** | 0.57 | 0.54 | 0.64 |
| UZ | 0.62 | **1.96** | 0.69 | **0.99** | 0.55 | 0.73 | 0.67 |
| OF.Var.Velo | 0.38 | **2.18** | **0.34** | 0.43 | 0.57 | 0.63 | 0.70 |
| AP.Dist | *-14.73* | 2.78 | 0.27 | 0.43 | 36.36 | ***0.94*** | **0.99** |
| AP.Lat.Mov | **5.20** | 0.02 | 0.27 | 0.10 | ***-5.44*** | 3.43 | 0.66 |
| AP.Var.Velo | -0.90 | 0.41 | 0.08 | 0.08 | ***2.09*** | -0.15 | 0.28 |
| Aguila (AZ) | | | | | | | |
| Latency | 20.67 | 0.60 | 0.57 | 0.54 | 0.76 | 0.68 | 0.61 |
| OF.Dist | -2.01 | 18.48 | **0.99** | **0.99** | 0.69 | 0.79 | 0.60 |
| UZ | -0.17 | **1.48** | 0.37 | **0.99** | 0.54 | 0.70 | 0.47 |
| OF.Var.Velo | -0.13 | **1.52** | **0.17** | 0.30 | 0.61 | *0.80* | 0.65 |
| AP.Dist | -4.28 | 2.80 | 0.10 | 0.21 | 17.89 | *0.83* | **0.99** |
| AP.Lat.Mov | 0.91 | 1.13 | 0.09 | *0.15* | *-1.27* | 1.61 | 0.69 |
| AP.Var.Velo | -0.19 | 0.15 | 0.00 | 0.03 | **0.91** | -0.06 | 0.16 |
| Socorro (NM) | | | | | | | |
| Latency | 20.57 | 0.54 | 0.65 | 0.54 | 0.51 | 0.55 | 0.46 |
| OF.Dist | -1.91 | 26.26 | **0.99** | **1.00** | **0.96** | ***0.90*** | ***0.91*** |
| UZ | -0.48 | **2.16** | 0.46 | **1.00** | ***0.91*** | *0.89* | *0.82* |
| OF.Var.Velo | -0.23 | **2.78** | **0.29** | 0.41 | ***0.94*** | ***0.92*** | ***0.92*** |
| AP.Dist | -0.74 | **12.83** | ***1.24*** | ***1.58*** | 24.39 | **0.99** | **0.99** |
| AP.Lat.Mov | 0.24 | ***-1.68*** | *-0.20* | ***-0.23*** | **-2.54** | 1.09 | 0.76 |
| AP.Var.Velo | 0.07 | ***0.97*** | *0.08* | ***0.13*** | **1.68** | -0.09 | 0.22 |
| Las Cruces (NM) | | | | | | | |
| Latency | 22.00 | 0.55 | 0.64 | 0.68 | 0.74 | 0.69 | 0.77 |
| OF.Dist | 0.43 | 12.46 | **0.99** | **1.00** | ***0.92*** | *0.83* | *0.84* |
| UZ | -0.17 | **0.65** | 0.21 | **0.99** | *0.86* | *0.84* | 0.77 |
| OF.Var.Velo | 0.26 | **1.02** | **0.09** | 0.18 | *0.87* | 0.76 | ***0.94*** |
| AP.Dist | 3.16 | ***4.90*** | *0.41* | *0.54* | 14.70 | **0.97** | **1.00** |
| AP.Lat.Mov | -0.59 | *-0.92* | *-0.10* | -0.09 | **-1.67** | 1.30 | 0.72 |
| AP.Var.Velo | 0.34 | *0.38* | 0.03 | ***0.07*** | **0.90** | -0.06 | 0.13 |

**Table S4.** Eigen decomposition of **G** for each population. Bold values indicate loadings > 0.25 to help interpretation.

| | $g_{max}$ | $g_2$ | $g_3$ | $g_4$ | $g_5$ | $g_6$ | $g_7$ | $g_{max}$ | $g_2$ | $g_3$ | $g_4$ | $g_5$ | $g_6$ | $g_7$ |
|---|---|---|---|---|---|---|---|---|---|---|---|---|---|---|
| | Dunnigan (CA) | | | | | | | Aguila (AZ) | | | | | | |
| λ | 51.11 | 24.59 | 17.64 | 2.28 | 0.52 | 0.17 | 0.13 | 25.47 | 17.59 | 14.52 | 1.38 | 0.27 | 0.16 | 0.11 |
| % Variance | 53.00 | 25.50 | 18.29 | 2.37 | 0.54 | 0.17 | 0.14 | 42.81 | 29.56 | 24.40 | 2.32 | 0.46 | 0.26 | 0.18 |
| Latency | **-0.66** | **-0.47** | **0.57** | -0.11 | 0.00 | 0.01 | -0.01 | **0.70** | **-0.53** | **0.48** | -0.04 | 0.00 | 0.00 | 0.00 |
| OF.Dist | 0.03 | **-0.78** | **-0.61** | -0.02 | -0.12 | -0.05 | -0.04 | **-0.44** | **-0.84** | **-0.29** | -0.08 | 0.11 | 0.03 | -0.01 |
| UZ | 0.00 | -0.08 | -0.04 | 0.12 | **0.92** | **-0.33** | -0.15 | -0.03 | -0.07 | -0.03 | -0.02 | **-0.88** | **0.42** | -0.20 |
| OF.Var.Velo | 0.00 | -0.09 | -0.05 | 0.05 | **0.34** | **0.72** | **0.59** | -0.03 | -0.07 | -0.02 | 0.04 | **-0.46** | **-0.79** | **0.40** |
| AP.Dist | **0.73** | **-0.40** | **0.54** | 0.10 | -0.03 | -0.04 | 0.05 | **-0.57** | 0.00 | **0.82** | 0.08 | -0.01 | 0.03 | 0.04 |
| AP.Lat.Mov | -0.16 | -0.01 | 0.00 | **0.98** | -0.14 | -0.05 | 0.05 | 0.04 | -0.09 | -0.07 | **0.99** | 0.01 | 0.04 | -0.02 |
| AP.Var.Velo | 0.04 | -0.03 | 0.02 | 0.08 | 0.07 | **0.60** | **-0.79** | -0.03 | 0.00 | 0.04 | 0.00 | 0.00 | **-0.45** | **-0.89** |
| | Socorro (NM) | | | | | | | Las Cruces (NM) | | | | | | |
| λ | 39.13 | 20.44 | 12.52 | 0.83 | 0.29 | 0.11 | 0.08 | 23.81 | 17.36 | 8.39 | 1.09 | 0.19 | 0.10 | 0.06 |
| % Variance | 53.31 | 27.85 | 17.06 | 1.13 | 0.39 | 0.14 | 0.11 | 46.69 | 34.05 | 16.46 | 2.13 | 0.37 | 0.19 | 0.11 |
| Latency | 0.10 | **0.99** | -0.10 | 0.01 | -0.02 | 0.01 | 0.00 | **0.86** | **0.49** | -0.13 | -0.01 | 0.01 | 0.02 | 0.00 |
| OF.Dist | **-0.72** | **0.01** | **-0.68** | -0.03 | 0.10 | 0.05 | 0.05 | 0.23 | **-0.61** | **-0.75** | -0.03 | -0.07 | 0.04 | 0.03 |
| UZ | -0.06 | -0.01 | -0.04 | 0.08 | **-0.95** | 0.19 | 0.20 | 0.01 | -0.04 | -0.03 | 0.04 | **0.91** | **0.37** | 0.18 |
| OF.Var.Velo | -0.08 | 0.00 | -0.06 | 0.04 | **-0.26** | **-0.63** | **-0.72** | 0.03 | -0.05 | -0.06 | -0.01 | **0.39** | **-0.67** | **-0.62** |
| AP.Dist | **-0.67** | 0.14 | **0.72** | -0.09 | 0.00 | 0.06 | -0.05 | **0.44** | **-0.62** | **0.64** | -0.10 | -0.02 | 0.04 | -0.04 |
| AP.Lat.Mov | 0.08 | -0.01 | -0.06 | **-0.98** | -0.09 | 0.07 | -0.09 | -0.07 | 0.08 | -0.04 | **-0.99** | 0.03 | 0.05 | -0.02 |
| AP.Var.Velo | -0.05 | 0.02 | 0.04 | -0.12 | -0.02 | **-0.74** | **0.66** | 0.03 | -0.04 | 0.03 | -0.05 | 0.11 | **-0.64** | **0.76** |

**Table S5**. Average temperature and rainfall at each population. Data from www.usclimatedata.com

| Site | Average summer max temperatures (C) | | | Average annual rainfall (mm) |
|---|---|---|---|---|
| | July | August | September | |
| Aguila (AZ) | 38.7 | 37.7 | 34.7 | 227.3 |
| Dunnigan (CA)* | 34.4 | 34.6 | 31.9 | 543.1 |
| Las Cruces (NM) | 34.9 | 33.4 | 30.9 | 247.9 |
| Socorro (NM) | 33.2 | 31.7 | 28.7 | 260.6 |

*data for Dunnigan, CA are from Woodland, CA

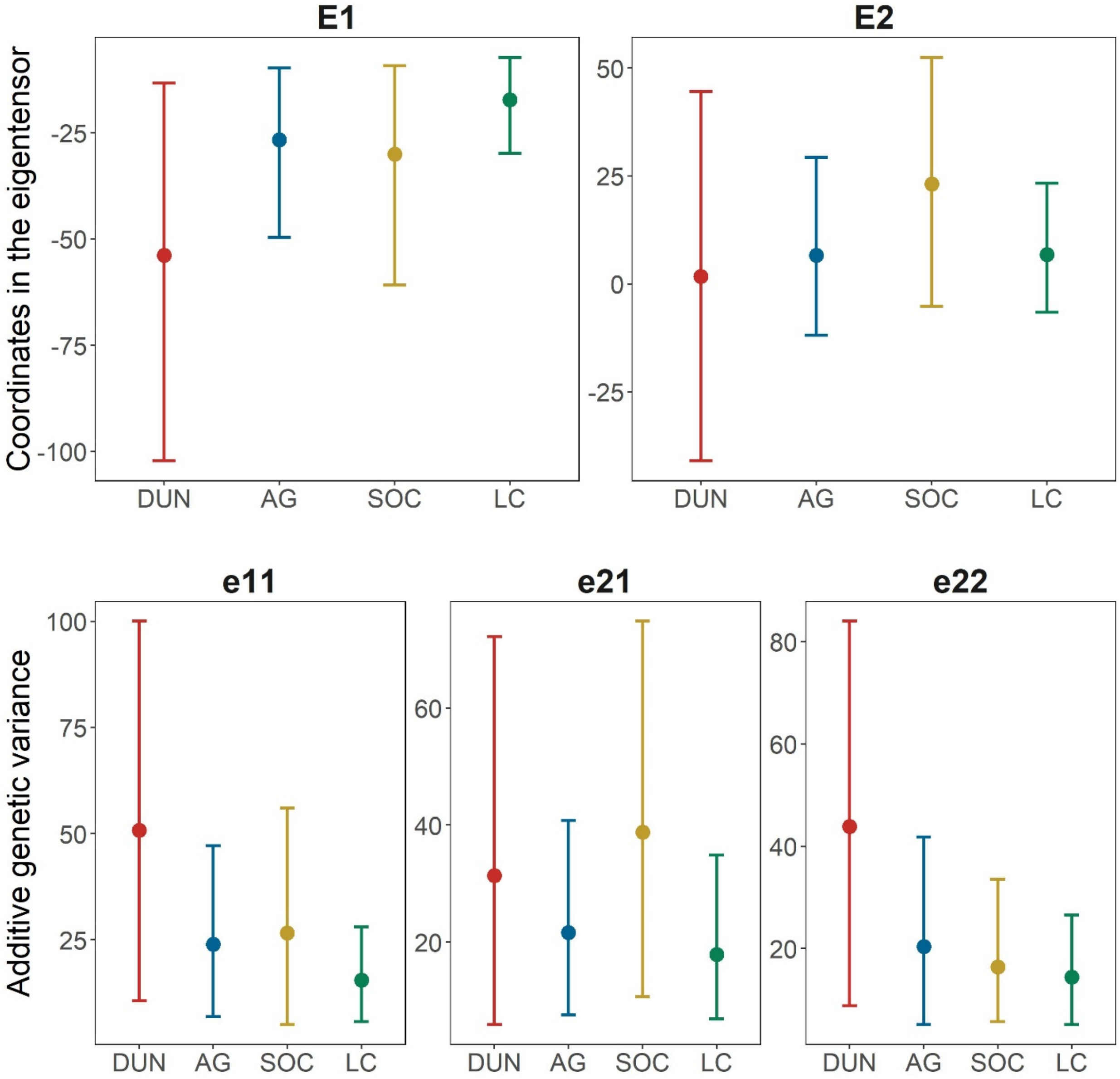


**Figure S1.** (a) Population coordinates along the leading eigentensors capturing most of the variation in divergence (**E**1 and **E**2). (b) Genetic variance in the direction of the leading eigenvectors of eigentensor **E**1 and **E**2. The largest population differences were found between the Dunnigan (DUN) and Las Cruces (LC) populations along **E**1 (Pmcmc > 0.85) and its leading eigenvector $\mathbf{e}_{11}$ (Pmcmc > 0.70), all other Pmcmc < 0.70.

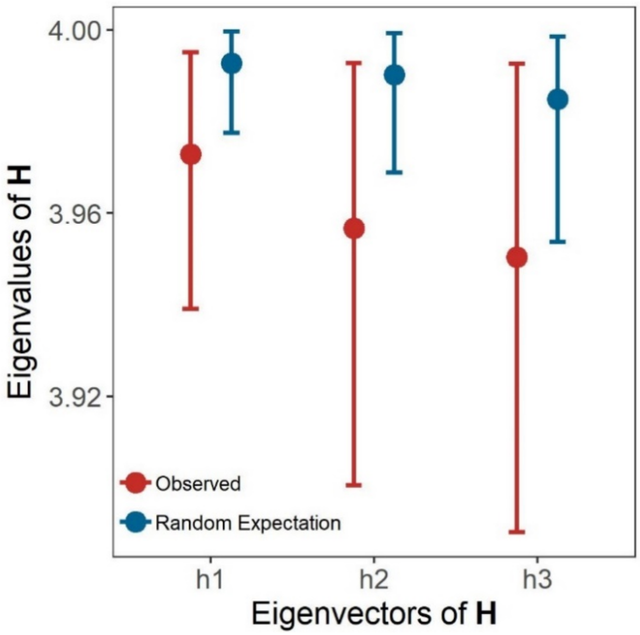


**Figure S2.** Eigenvalues of **H** for a comparison of the first three eigenvectors of each **G** matrix compared against random expectations. None of the eigenvectors of **H** showed significant departure from random expectations (Pmcmc < 0.65), indicating that major axes of genetic variation are highly conserved among populations. (h1: Pmcmc = 0.6, h2: Pmcmc = 0.65, h3: Pmcmc = 0.65)

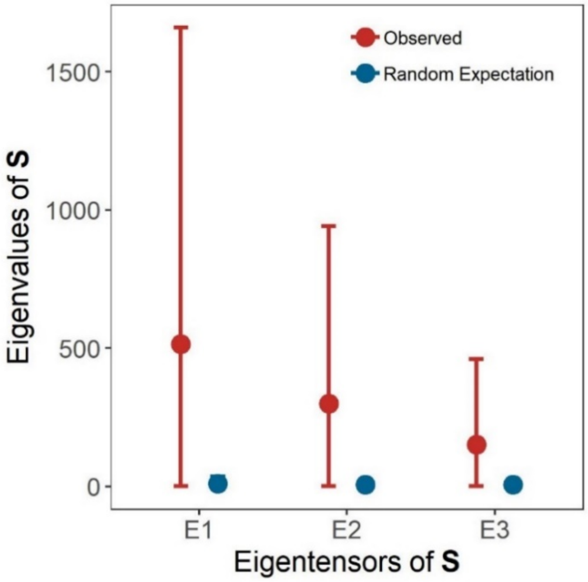


**Figure S3.** Eigenvalues of the first three positive eigentensors of **S**. Both $\mathbf{E}_1$, $\mathbf{E}_2$ and $\mathbf{E}_3$ were significantly different than expected from the random distribution (Pmcmc > 0.85), indicating that populations differed along $\mathbf{E}_1$, $\mathbf{E}_2$ and $\mathbf{E}_3$. (E1: Pmcmc = 0.85, E2: Pmcmc = 0. 85, E3: Pmcmc = 0. 85)
